# Maternal diet and gut microbiome composition modulate early life immune development

**DOI:** 10.1101/2023.03.06.531289

**Authors:** Erica T Grant, Marie Boudaud, Arnaud Muller, Andrew J Macpherson, Mahesh S Desai

## Abstract

In early life, the intestinal mucosa and immune system undergo a critical developmental process to contain the expanding gut microbiome while promoting tolerance towards commensals, yet the influence of maternal diet and gut microbial composition on offspring immune maturation remains poorly understood. We colonized gnotobiotic mice with a defined consortium of 14 strains, fed them a standard fiber-rich chow or a fiber-free diet, and then longitudinally assessed offspring development during the weaning period. Unlike pups born to dams fed the fiber-rich diet, pups of fiber-deprived dams demonstrated delayed colonization with *Akkermansia muciniphila*, a mucin-foraging bacterium that can also utilize milk oligosaccharides. The pups of fiber-deprived dams exhibited an enrichment of colonic tissue transcripts corresponding to defense response pathways and a peak in *Il22* expression at weaning. Removal of *A*. *muciniphila* from the community, but maintenance on the fiber-rich diet, was associated with reduced proportions of RORγt-positive innate and adaptive immune cell subsets. Our results highlight the potent influence of maternal dietary fiber intake and discrete changes in microbial composition on the postnatal microbiome assemblage and early immune development.

## Introduction

Dietary fiber plays a major role in shaping the colonic gut microbiome and host immunity, however the impact of maternal dietary fiber intake on the offspring’s microbiome and immune development is largely associative for the early life period (Mirpuri, 2021). In adult mice, dietary fiber deprivation leads to microbiome-driven thinning of the mucus layer, which can increase susceptibility to a range of conditions including enteric pathogen infection (Desai *et al*., 2016; Neumann *et al*., 2021), graftversus-host disease (Hayase *et al*., 2022), and food allergy (Parrish, Boudaud, *et al*., 2022). Although exposure to microbial metabolites in utero initiates innate immune responses and barrier development in pups (De Agüero *et al*., 2016), the postnatal period is also critical for healthy maturation of intestinal function and adaptive immunity in a complex process that is partially driven by a response to the colonizing microbiome (Kalbermatter *et al*., 2021; Wells *et al*., 2022). In particular, from 2–4 weeks of age, mouse pups transition from breast milk to solid food with an accompanying shift in microbial composition and a vigorous immune response to the expanding microbiota (Al Nabhani *et al*., 2019). Without this “weaning reaction”, as observed in germ-free and broad-spectrum antibiotic-treated mice, Al Nabhani *et al* reported increased susceptibility to inflammatory pathologies (Al Nabhani *et al*., 2019). However, it remains unclear how variations in the maternal diet or defined differences in microbial composition affect early life immune profile (Ansaldo *et al*., 2019).

We therefore set out to investigate the influence of maternal dietary fiber intake on the offspring colonic development by feeding dams either a standard fiber-rich (FR) chow or a custom fiber-free (FF) diet (Desai *et al*., 2016) (**Fig 1A**). To standardize the bacterial composition and facilitate interpretation of the microbial shifts, we utilized a gnotobiotic mouse model consisting of 14 bacterial strains (14-member synthetic human gut microbiota or 14SM) that form a functionally diverse community (Desai *et al*., 2016; Steimle *et al*., 2021). Although this community was designed to resemble an adult microbiota, one member—*Akkermansia muciniphila—*is capable of degrading milk oligosaccharides (Kostopoulos *et al*., 2020; Luna *et al*., 2022) and is an early colonizer of the gut (Collado *et al*., 2007). This taxon is also enriched under fiber-deprived conditions in adult mice (Desai *et al*., 2016) and appears to have immunogenic properties when present in 14SM, including a high IgA-coating index (Martens *et al*., 2023). We sought to determine the role of this particular microbe in early life development on the same FR diet by employing mice colonized with the full community except *A. muciniphila* (13SM). By employing a longitudinal sampling approach among the pups of each diet and SM combination, we isolated dietary and microbial drivers of early-life immune development. We show that both the maternal diet and the presence or absence of *A. muciniphila* are key contributing factors in shaping the host colonic transcriptome and the immune cell populations during this critical early life window.

**Figure 1.**
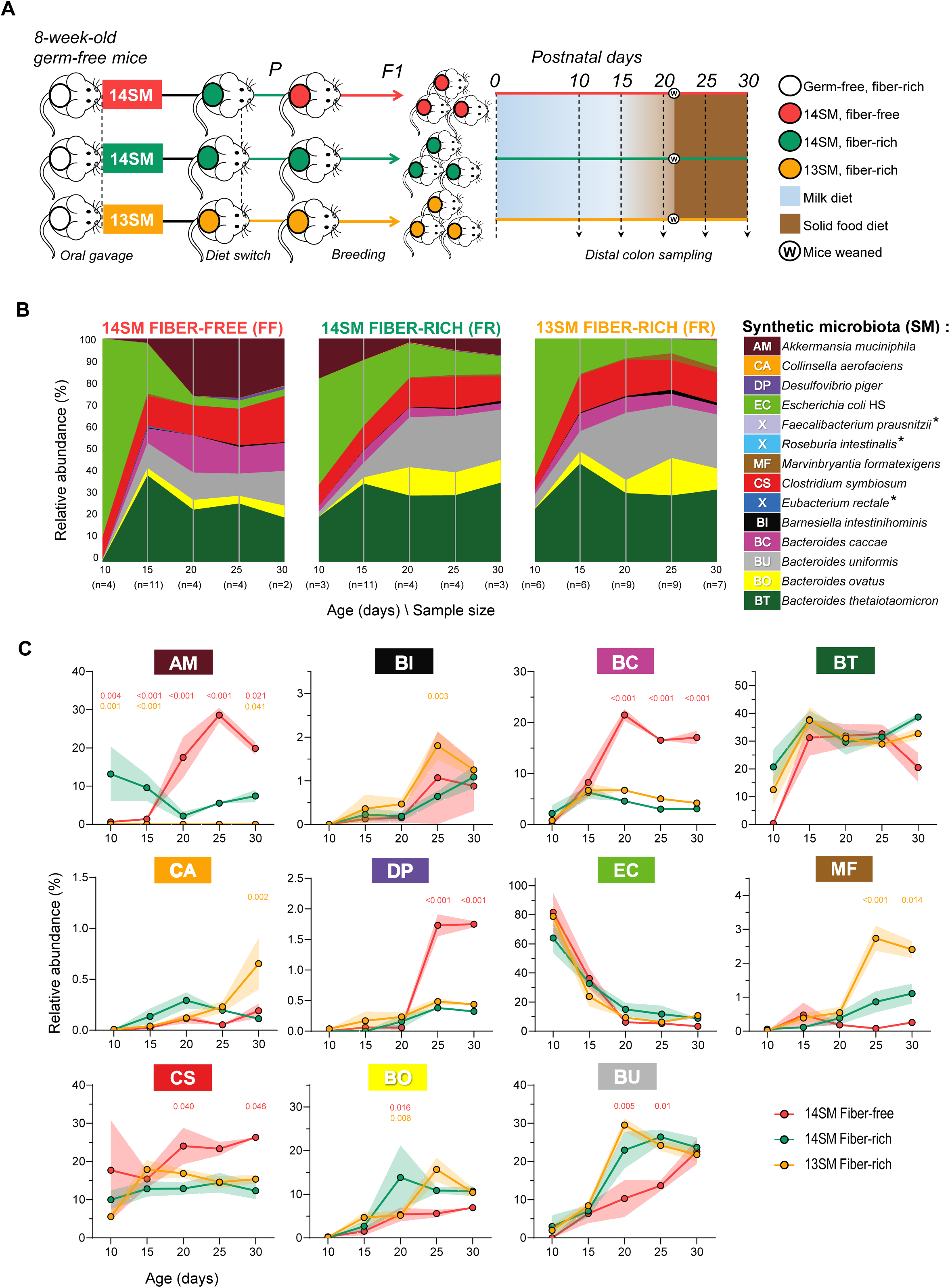
Delayed expansion of *Akkermansia* among offspring of fiber-deprived dams. **A**. Schematic representation of the study design. **B**. Stream plot showing the relative abundance of the 14 bacterial strains in the colons of pups born to 14SM FF-fed mice, 14SM FR-fed mice and 13SM FR-fed dams. Abundances represent an average of all pups at the specified age. Distal colon contents were analyzed by qPCR using primers against targets that are unique to each bacteria and present once per genome. Asterisk indicates bacterial strains that were considered absent (average abundance <0.1%) in the pups. **C**. Line plots of the mean relative abundances and SD of each bacterial strain from ages 10 to 30 days. For non-AM strains, relative abundances of the 14SM groups were recalculated excluding AM in order to compare abundances with the 13SM group. Data information: Strain abundances were normally distributed within each group according to the Kolmogorov-Smirnov test. Statistical significance compared to the 14SM FR control at each time point was calculated by two-way ANOVA with false-discovery adjustment using the Benjamini and Hochberg method. Adjusted *p* values are indicated for 14SM FF vs 14SM FR (red) and 13SM FR vs 14SM FR (orange). Data are from two independent experiments and represent biological replicates (*n* = 2–11/group).

## Results and discussion

### Gut microbiome colonization is shaped by maternal diet

Consistent with previous reports (Earle *et al*., 2015; Desai *et al*., 2016), we observed an elevated relative abundance of *Akkermansia muciniphila* in the parent mice fed a fiber-free (FF) diet compared to those on a standard fiber-rich (FR) chow (**Dataset EV1**). We expected to observe similarly high levels of *A. muciniphila* in the offspring of FF-fed dams; however, the expansion of *A. muciniphila* was delayed among mice born to FF-fed dams (**Figs 1B–C**). This contrasts with the microbiome development of 14SM FR pups, which showed strong initial colonization with *A. muciniphila* during suckling, as anticipated given its ability to consume milk oligosaccharides (Kostopoulos *et al*., 2020). When mice began to consume a solid food diet between ages 15 to 20 days, the profiles rapidly stabilized and resembled compositions observed in adult mice fed the respective diets (**Datasets EV1–2**), as also verified by 16S rRNA gene sequencing (**Dataset EV3**). Among 14SM FF mice, *A. muciniphila*, *Bacteroides caccae, Clostridium symbiosum*, and *Desulfovibrio piger* significantly expanded to comprise approximately three-fourths of the microbiome composition. As *B. caccae* can degrade complex polysaccharides and mucin, its elevated abundance on the FF diet may reflect a functional shift towards increased mucin foraging, which can ultimately contribute to thinning of the mucus barrier (Desai *et al*., 2016) and promote production of immunostimulatory cytokines TNF and IL-6 (Brown *et al*., 2021). Although the consequences of an increase in *C. symbiosum* are less clear from a functional standpoint, it is elevated in IBD and is an emerging biomarker for colorectal cancer (Xie *et al*., 2017). Similarly, *D. piger* may also contribute to development of local inflammation and colorectal cancer via production of hydrogen sulfide (Blachier *et al*., 2021), particularly when combined with increased liberation of sulfate moieties by mucin-foraging bacteria under fiber-free conditions.

Differences in the microbial assemblage between pups of 14SM and 13SM dams were less pronounced than between pups from FR and FF-fed dams. However, we note a transient increase in *Barnesiella intestinihominis*, a mucin-specialist, at day 25; *Collinsella aerofacians* at day 30, a bacterium linked to increased gut permeability; and decrease in *Bacteroides ovatus* at day 20, a bacterium linked to production of short-chain fatty acids and neuroactive compounds (Horvath *et al*., 2022). After weaning, there was also a marked increase in *Marvinbryantia formatexigens*, an oligo-saccharide degrading bacterium (Rey *et al*., 2010) that can stimulate host IgA production via acetate (Takeuchi *et al*., 2021). Although preliminary, these data indicate that the absence of *A. muciniphila* may have a distinct impact on the community structure and host immunity through diet-independent mechanisms.

### Colonic transcriptome reflects differences in maternal diet

*A. muciniphila* has been previously associated with regulation of host metabolic activity (Yoon *et al*., 2021) and intestinal adaptive immunity in adult mice (Ansaldo *et al*., 2019), but little is known about its immune influence in early life. Therefore, to address the host physiological effects of this striking difference in the abundance of *A. muciniphila* according to diet (**Fig 1**), we performed total RNA sequencing on the distal colon tissues of 15 day-old pups born to FR and FF 14SM dams as well as FR 13SM dams without *A. muciniphila* (**Fig 2, Fig EV1A, Dataset EV4**). Relative to the control group (14SM FR), we detected changes in the expression level of 32 transcripts according to the maternal diet (14SM FF) and 57 transcripts regulated by the presence of *A. muciniphila* (13SM FR) (**Fig 2A top, Fig 2B, Dataset EV5–6**). Performing a gene set analysis based on Gene Ontology (GO) terms for biological processes, we found differential enrichment according to the diet (14SM FR vs 14SM FF), but not to the presence of *A. muciniphila* (14SM FR vs 13SM FR) (**Fig 2A bottom, Fig 2C**). This may reflect two limitations of gene set analyses: 1) transcripts not corresponding to a common pathway can still directly lead to changes in immune cell differentiation; and 2) transcripts corresponding to pseudogenes or ncRNA may impact on host development through epigenetic mechanisms or other undescribed pathways (**Fig EV1B, Dataset EV6**). Such cases would not be highlighted in gene set analyses, therefore the absence of significant changes on the level of gene sets for the 14SM FR vs 13SM FR comparison does not mean there are no biologically relevant differences in the transcriptomes. For example, H2-K1 (a histo-compatibility gene involved in antigen presentation), Gapdh (involved in glycolysis), and Gal3st2b (involved in glycosylation) represent relevant differences between the 13SM and 14SM groups that can underlie the observed immunological differences, but which do not correspond to a shared functional pathway (**Dataset EV6**).

**Figure 2.**
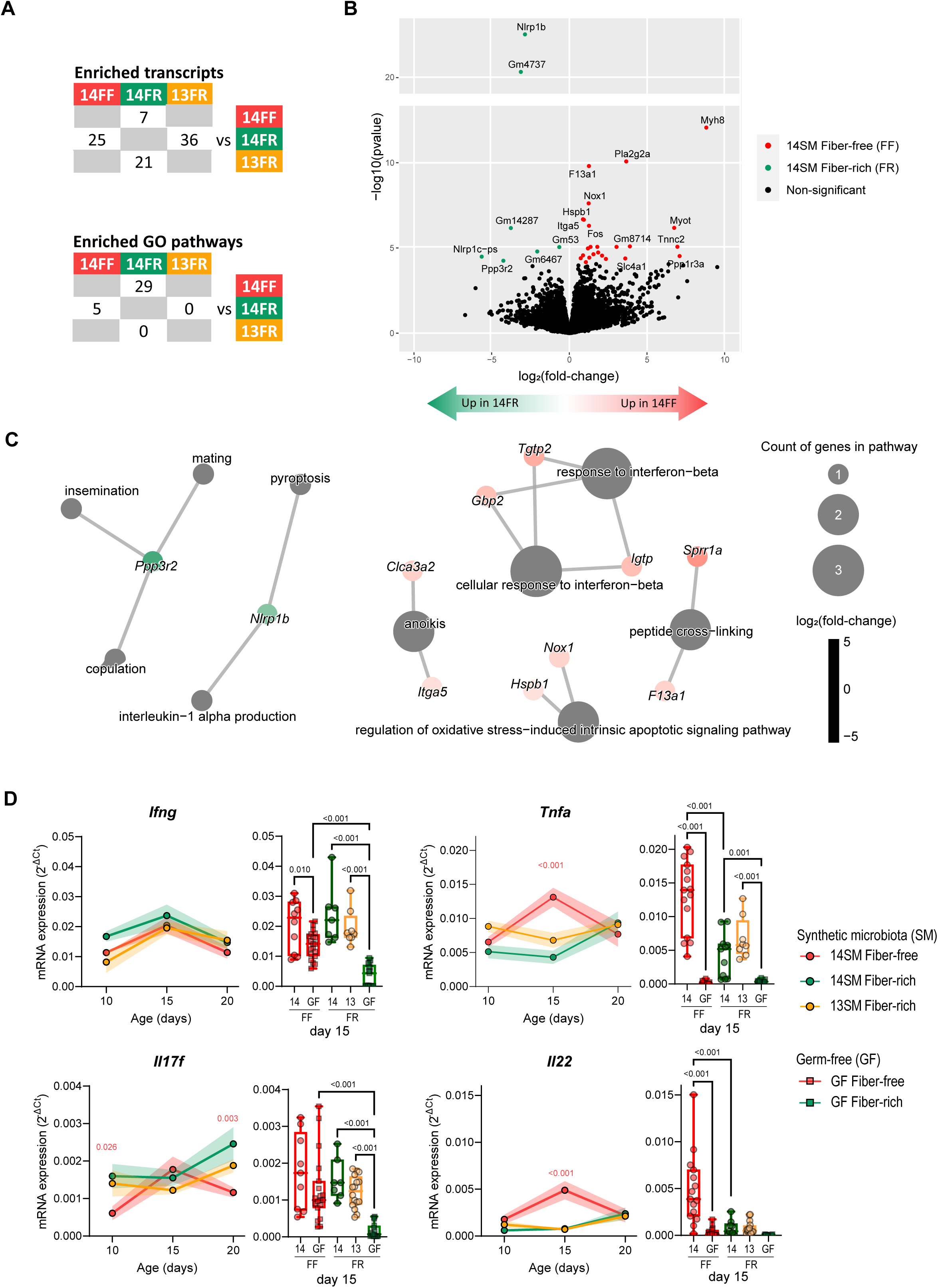
Maternal diet alters colonic transcriptome of offspring at age 15 days. **A**. Table depicting counts of genes with differential transcription across groups after filtering reads not present at least once across all samples (top; see **Dataset EV5, Dataset EV6**) and differentially enriched gene sets between groups according to the Gene Ontologies (GO) terms for biological processes (bottom; see **Dataset EV7**). **B**. Volcano plot showing gene abbreviations for colonic transcripts that were significantly enriched in 14SM FF or 14SM FR after correction for multiple comparisons using DESeq2 (see **Dataset EV5**). **C**. Category netplots displaying gene sets for biological processes and their associated genes that were differentially represented in pups born to 14SM FF vs 14SM FR dams. Top five up-and down-regulated gene sets for biological processes are shown; all gene sets can be found in **Dataset EV7**. **D**. mRNA expression of *Ifng*, *Tnfa*, *Il17f*, and *Il22*. Colonic transcription levels were normalized by *Hprt*, which was the most stable among the three housekeeping genes tested (*Hprt, Hsp90, Gapdh)*. Longitudinal data of mean ± SEM is shown for 14SM FF, 14SM FR, and 13SM FR pups at age 10-20 days (circles) with statistical significance calculated using two-way ANOVA factored on group and time point compared to the 14SM FR control. To the right of each longitudinal plot, we also report colonic transcript levels for pups born to germ-free (GF) FF or FR dams at age 15 days (squares) with statistical significance based on a one-way ANOVA with false-discovery adjustment using the Benjamini and Hochberg method. Outliers removed using ROUT method with Q = 10%. Data are from two independent experiments and represent biological replicates (*n* = 3–20/group).

On a functional level, *Ppp3r2* and *Nlrp1b* led to the identification of the top 5 gene sets enriched in 14SM FR pups (**Fig 2C;** see **Dataset EV7** for all 29 gene sets), though less weight should be placed on this finding since it was driven by only two genes. Comparing host transcriptomes according to diet revealed an enrichment in transcripts involved in epithelial barrier development and cell–cell adhesion (*Myh1*, *Myh8*, and *Myot*), as well as peptide cross-linking (*Sprr1a* and *F13a1*), anoikis or apoptosis associated with cell detachment from the epithelial tissue (*Clca3a2* and *Itga5*), oxidative stress-induced intrinsic apoptotic signaling pathways (*Hspb1* and *Nox1*), and response to interferon-beta (*Tgtp2*, *Gbp2* and *Igtp*) among FF 14SM pups (**Fig 2B–C, Table EV7**). Some of these transcripts, including *Nox1* (Szanto *et al*., 2005), *Itga5* (Zhu *et al*., 2021), and *Hspb1* (Arrigo *et al*., 2007), are also implicated in colon cancer progression, suggesting potential pathological development. The differential expression of these transcripts according to RNA sequencing was further validated using primers specific to these transcripts. The RT-qPCR results exhibited a trend consistent with expectations from RNA sequencing analyses, although these shifts were only significant for *Nox1*, *Ppp3r2*, and *Ifit1bl1* (**Fig EV1D–E**). Several transcripts belonging to the “response to interferon-beta” gene set were elevated among 14SM FF pups, however we detected only a slight increase in *Ifnb1* among this group at 15 days of age that was not maintained at other timepoints (**Fig EV1E**).

The postnatal period is critical for the development of the immune system as a weaning reaction to colonizing microbes was shown to be protective against diseases in adulthood among mice colonized with segmented filamentous bacteria (Al Nabhani *et al*., 2019). Thus, in order to provide a longitudinal element to the transcriptomic data, we performed RT-qPCR to quantify the relative expression of transcripts corresponding to select cytokines involved in immunity and epithelial barrier maintenance (**Fig 2D**). We noted an induction of *Infg* around 15 days among all groups, which is considered characteristic of the weaning reaction. *Tnfa*, another marker of the weaning reaction, was only elevated in the pups of 14SM fiber-deprived dams. By comparing transcript expression levels to germ-free (GF) mice at age 15 days (**Fig 2D**), we see that some of the observed pro-inflammatory shifts can be attributed to a combination of diet and microbiome composition. In particular, at age 15 days, the FF diet increased the transcript levels of *Ifng*, *Il17f*, and *Il23* among germ-free pups of fiber-deprived dams, relative to germ-free, fiber-rich-fed offspring (**Fig 2D, Fig EV1C**). Although the expression of these cytokines is overall lower in germ-free conditions compared to 13SM or 14SM, differences in the maternal diet may affect the baseline expression of these cytokines in germ-free mice through a possibly different maternal milk composition. At the same age, only pups of 14SM FF-fed dams displayed a strong elevation in *Tnfa* and *Il22* transcript levels, reflecting compounding effects of the maternal diet and microbiome.

### Maternal fiber deprivation shapes microbiota-mediated immune development

The transcript-level results underscore the potent impact of the maternal diet in shaping the developing epithelial barrier and host immune system. Therefore, to confirm that these changes translate into the profile of immune cell populations, we performed flow cytometry on cells isolated from the colonic lamina propria (cLP) at age 15 days (**Fig 3, Fig EV2**). As an effect of colonization, we observed elevated levels of cytotoxic NK cells and ILC3 among 14SM pups compared to GF pups (**Fig 3A**), in agreement with previous reports comparing germ-free and colonized mice (Ganal *et al*., 2012; Sanos *et al*., 2009). Consistent with the host transcriptomic data, maternal dietary fiber intake appeared to play a role in immune cell profiles of both 14SM and GF pups. Particularly, pups of 14SM FF dams had lower levels of ILC2 and higher levels of NCR^-^ ILC3 compared to pups of 14SM FR dams (**Fig 3A**). This aligns with observations by Babu et al. among pups of high-fat diet-fed mothers, suggesting common underlying mechanisms driven by a Western-style diet (Babu *et al*., 2018). In the adaptive arm however, the 14SM colonization lowered the proportions of CD8^+^ T cells, CD4^+^ T cells, Tregs and Th1 cells (**Fig 3B**). By contrast, Th17 cells were induced by the colonization among pups of FR-fed dams but not fiber-deprived dams, suggesting that the maternal dietary fiber intake differentially affects innate and adaptive RORgt^+^ cells in the pup’s colons.

**Figure 3.**
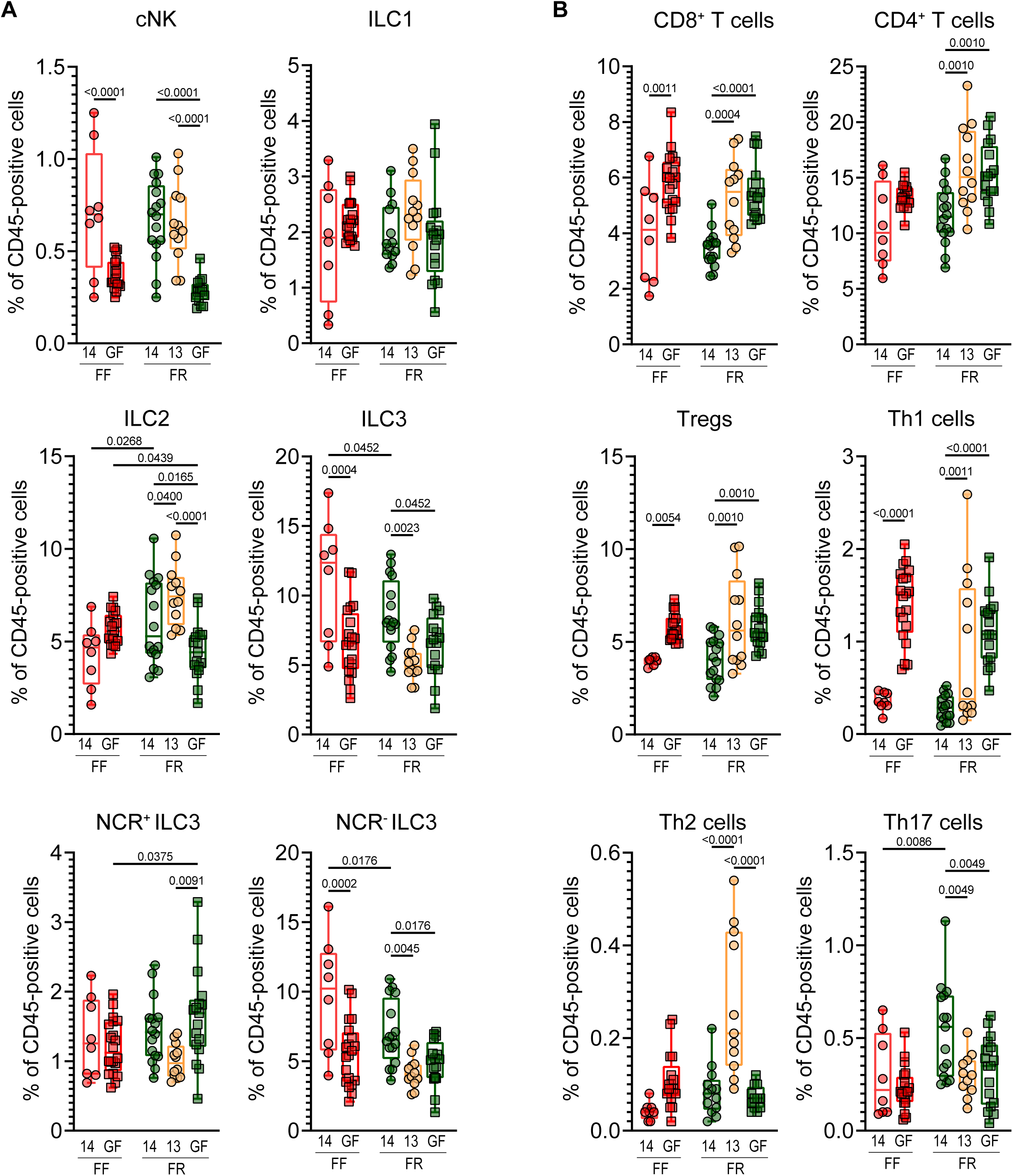
Maternal diet and microbiome shape innate and adaptive neonatal immune cell populations. **A**. For mice at age 15 days, the proportion of cytotoxic natural killer (cNK), innate lymphoid cell (ILC) type 1, ILC type 2, ILC type 3, natural cytotoxicity receptor (NCR) positive ILC3, and NCR negative ILC3 populations among CD45^+^ cells. **B**. For mice at age 15 days, the proportion of CD8^+^ T cells, CD4^+^ T cells, Tregs (total FoxP3^+^), Th1 cells (T-bet^+^FoxP3^-^), Th2 (GATA3^+^FoxP3^-^), and Th17 (RORgt^+^FoxP3^-^) populations among CD45^+^ cells. Statistical significance is based on a one-way ANOVA with false-discovery adjustment using the Benjamini and Hochberg method. Data are from two independent experiments and represent biological replicates (*n* = 8–20/group).

Bacteria-derived metabolites represent a mechanistic link between changes in the microbiome and host immunity. While it would be valuable to perform metabolomics analysis during this early life period, we note that the overall volume of colonic contents available is a major limitation, especially at 10–15 days of age. In the absence of such data, we would speculate that microbial metabolism of tryptophan, which is abundant in milk, may drive many of the observed microbiota-mediated immune changes. Microbial or host degradation of dietary tryptophan yields indole derivatives which can function as AhR ligands and stimulate production of IL-22 from ILC3s (Qiu *et al*., 2012; Li *et al*., 2018), both of which were elevated among pups of 14SM FF-fed dams at 15 days of age (**Fig 2D, 3A**). The expansion of these innate subsets also leads to apoptosis of commensal-specific CD4+ and CD8+ T cells (Hepworth *et al*., 2015; Liu *et al*., 2019). Likewise, among adaptive cell subsets, Th17 cells were significantly elevated among 14SM FR mice, potentially reflecting the effects of increased colonization with *A. muciniphila* in early life, since this bacterium can act as a direct AhR ligand via its outer membrane protein Amuc_1100 (Gu *et al*., 2021). Finally, we note that a fixed number of cells were stained per organ, therefore the higher proportions of adaptive cell subsets (namely CD8^+^, CD4^+^, Tregs, and Th1 cells) observed in GF mice may simply reflect a lower absolute number of innate cells (Macpherson and Harris, 2004; Kennedy *et al*., 2018).

### Akkermansia muciniphila alters innate and adaptive immunity in early life

To determine the role of *A. muciniphila* in the aforementioned colonization effect, we compared the immune cell profiles of pups born to FR-fed 14SM, 13SM and GF dams (**Fig 3**). Under FR conditions, colonization differentially affected the proportions of innate cell subsets, with ILC3s favored under 14SM conditions (p=0.0023) versus ILC2 cells under 13SM conditions (p=0.0400) (**Fig 3A**). Although the FF diet skewed toward ILC2 in the absence of a microbiota, the strong enrichment of ILC3s in 14SM colonized mice (p=0.004) led to a null effect when comparing ILC2 proportions in pups of 14SM and GF FF-fed dams directly. These results highlight the complex interactions between diet and microbiota on immune development in early life.

Interestingly, an increase of NCR^-^ ILC3 and Th17, and the decrease of CD8^+^ T cells, CD4^+^ T cells, Tregs and Th1 cells in pups born to 14SM FR-fed dams, relative to those of GF FR-fed dams, was not observed in pups born to 13SM dams, confirming a specific impact of *A. muciniphila* on these immune cell populations. Furthermore, only Th17 cells were decreased in pups born to 14SM FF-fed dams, consistent with a reduced relative abundance of *A. muciniphila* at this time point compared to pups born to 14SM FF-fed dams (**Fig 3B**). These results support an early life induction of the colonic Th17 population that is driven by the maternal dietary fiber intake and mediated locally by *A. muciniphila* in the pups colons. Th17 cells play a key role in progression of multiple sclerosis, an autoimmune disease that has also been correlatively linked to presence of *A. muciniphila* in patients (Jangi *et al*., 2016) and mechanistically in mouse models of multiple sclerosis (Lin *et al*., 2021). Although the precise nature of the link between *Akkermansia* and exacerbation of autoimmune encephalomyelitis has not yet been shown (e.g. MAMP recognition, physiological alterations from mucin-degrading activities), this relationship bears further investigation to identify potential therapeutic targets.

## Conclusion

Summarizing these findings, we demonstrate: 1) that the maternal diet alters the host colonic transcriptomic and immune profile among pups during the weaning period; and 2) that these observed alterations can also be tuned by the inclusion or exclusion of a single bacterium from a defined community (**Fig 4**). The mechanisms behind such immunomodulatory effects can be via microbial metabolites such as SCFA or AhR ligands, or via surface molecules such as LPS (Davis *et al*., 2022). In the latter instance, Kostopoulos et al. demonstrated that *A. muciniphila* produces more immune-regulatory pili-protein when grown on human milk, highlighting diet-dependent factors that can indirectly strengthen barrier integrity via the microbiome (Kostopoulos *et al*., 2020). In line with this, we found that presence of *A. muciniphila* skews toward a type 3 response, whereas its absence appears to favor a type 1 or 2 response. Overall, our results highlight the importance of the maternal microbiome composition and dietary fiber intake in postnatal microbiome maturation and immune development, which could have implications for development of various immune-mediated diseases later in life.

**Figure 4.**
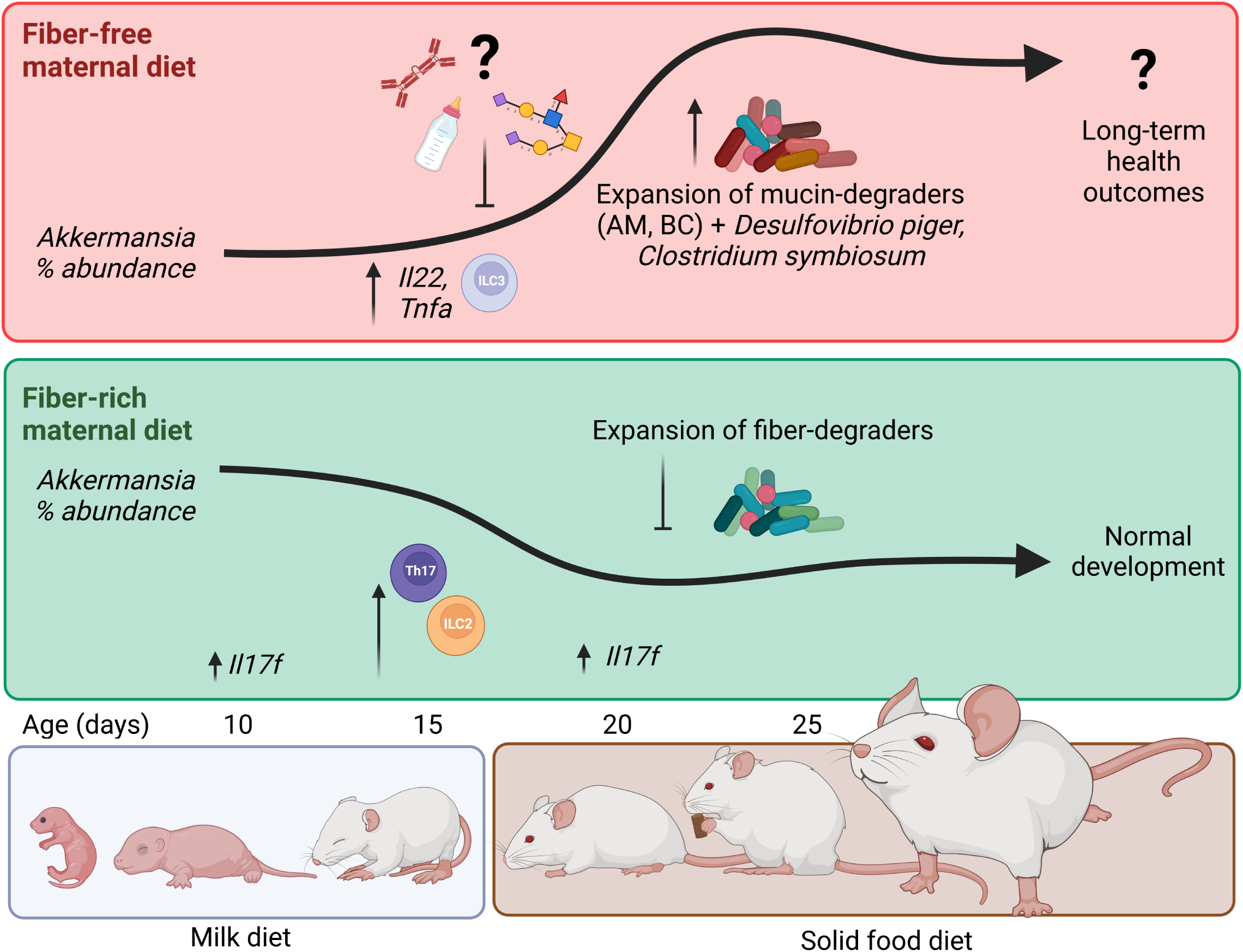
Summary of key transcriptional, cellular, and microbial shifts among pups of 14SM fiber-deprived dams. Maternal milk components (immunoglobulins, altered milk glycan structures) among fiber-deprived dams are predicted to inhibit the establishment of mucin-degrading bacteria in the suckling period, after which this functional group rapidly expands, following a pattern inverse to pups of FR-fed dams. These microbial changes are accompanied by an array of host changes characterized by colonic transcripts involved in epithelial development and innate defense signaling as well as differential development of key immune cell subsets.

In this study, we assessed the compounding effects of a fiber-free diet across generations with mice pups weaned onto the same diet as the dams. While the observed immune changes occur prior to weaning, modifying the diet at distinct developmental stages (in utero, pre-weaning, and post weaning) would provide insights into critical key drivers of early life immune development. Furthermore, it would be of interest to interrogate both the persistence of the observed differences in early life immunity in an adult mouse and the effects of this differential development in a relevant disease model (**Fig 4**). For example, Al Nababi et al. showed that the weaning reaction provided protection against oxazolone-induced colitis—a type-2 inflammation—and, considering the enrichment type 2 immune cell subsets in pups of 13SM FR-fed mice at 15 days of age, this could be an promising model to further investigate diet and microbiota effects on immunity at distinct developmental stages. Thorburn et al., for example, found that fiber supplementation during pregnancy alone was sufficient to prevent asthma development in offspring (Thorburn *et al*., 2015). Although the involvement of the microbiome was not explored by Thorburn et al., we previously documented the synergistic impact of diet and *A. muciniphila* in the context of food allergy (Parrish, Boudaud, *et al*., 2022). However, the study by Parrish et al. was performed in adult mice that were colonized by oral gavage, therefore the question remains as to how exposure to *A. muciniphila* in early life might play into the development and severity of allergic responses. Finally, mouse models of multiple sclerosis might also be of interest considering RORgt+ immune cell subsets were strongly affected by the maternal diet or microbiome changes in the present study and this same subset is also known to drive the demyelination that characterizes this disease.

Additionally, future studies should characterize the maternal milk components to improve understanding of the mechanisms for *A. muciniphila* suppression, which occurred only in pups born to fiber-deprived dams. As illustrated in **Fig 4**, a potential explanation for this phenomenon includes changes in quantity or specificity of IgG—the dominant maternal antibody transferred via murine breast milk (Zheng *et al*., 2020)—which can be also specifically induced by *A. muciniphila* (Ansaldo *et al*., 2019). Follow-up studies should be performed to investigate whether increased *A. muciniphila* and mucus erosion among FF-fed dams facilitates immune recognition of this bacterium, leading to increased the production of specific-IgG in breastmilk, ultimately preventing *A. muciniphila* colonization and Th17 induction among offspring in early life. Another explanation for *A. muciniphila* suppression may be related to differences in the types of linkages presented by the milk oligosaccharides, which could hamper their effective utilization by *A. muciniphila.* To this point, Kostopoulos et al. demonstrated that *A. muciniphila* can degrade 2’-fucosyllactose, lacto-N-tetraose, lacto-N-triose II, lactose, and 3’-sialyllactoses (Kostopoulos *et al*., 2020), although it also bears mentioning that these structures are common components of human milk, which is expected to have a slightly different compositional and structural makeup compared to murine milk (Luna *et al*., 2022). Improved understanding of these milk-associated mechanisms in both humans and rodent models are warranted, as these insights could provide tools to augment growth or activity of *A. muciniphila* in various disease contexts.

While the 14SM model enables controlled investigation of the effects the microbiome on host immune development, it is important to note that the 14SM consortium is based on an adult human colonic microbiome. Although the species in this community are closely related to strains that are normally found in the mouse gut and thus readily colonise upon oral gavage in germ-free mice (Desai *et al*., 2016), the 14SM lacks certain members, such as bifidobacteria and lactobacilli, which are abundant in early life and mostly colonize the small intestine. These members could be added to the community in future studies, however we would also highlight that a community strictly designed to model the pre-weaning mouse gut would present its own set of limitations. To this point, Lubin et al. demonstrated that restricting the microbiome to a pre-weaning community halts immune development and increases susceptibility to *Salmonella* infection (Lubin *et al*., 2023), reflecting the need for a functionally-balanced synthetic community to elicit normal immune development. As with other gnotobiotic models, the 14SM immune profiles typically represent an intermediate between germ-free and SPF colonization states in adulthood, potentially due to low diversity, low bacterial density and/or absence of specific immunogenic bacteria such as segmented fillamentous bacteria (Parrish, Grant, *et al*., 2022). Addition of new members may be a promising approach to refine these models, although it remains difficult to completely recapitulate SPF immune profiles (Afrizal *et al*., 2022). Despite these limitations, synthetic microbiota models remain critical tools to dissect specific interactions between the microbiome and diet in early life.

This work highlights numerous underexplored factors related to the potential for maternal fiber deprivation to alter postnatal establishment of mucin-degrading gut bacteria and immune maturation. The elevated levels of RORgt-induced innate and adaptive cell subsets among *A. muciniphila*-colonized mice is important for preventing translocation of gut microbes into the host (De Agüero *et al*., 2016) and can also be associated with protection against chronic inflammation via TLR4 (Liu *et al*., 2022). Considering *A. muciniphila* is only found in approximately half of the human population (Geerlings *et al*., 2018), linking this bacterium to exacerbation or amelioration of specific gut-linked diseases is particularly interesting as it highlights its potential as a biomarker for disease risk. Although *A. muciniphila* has been proposed as a probiotic (Cheng and Xie, 2021), mainly due to its strong benefits in countering metabolic diseases, the various conflicting reports involving this bacterium and health, as discussed by Cirstea et al. (Cirstea *et al*., 2018), caution against premature generalizations and underscore the need to consider factors such as diet and underlying disease context. Ultimately, insights provided by the present study and follow-up works are expected to hold translational relevance in improving human health outcomes as modern lifestyle practices (e.g. antibiotics use, Western-style diets) contribute to perturbation of the gut microbiota and immune function across generations (Sonnenburg *et al*., 2016) and in early life (Robertson *et al*., 2019).

## Materials and Methods

### Mice

All experimental procedures involving animals (i.e. initial gavage of parental mice) were approved by the Luxembourgish Ministry of Agriculture, Viticulture and Rural Development (LUPA 2019/49). Mouse work was performed according to the “Règlement grand-ducal du 11 janvier 2013 relatif à la protection des animaux utilisés à des fins scientifiques” based on the “Directive 2010/63/EU” on the protection of animals used for scientific purposes. Mice were housed in the germ-free facility of the University of Luxembourg in either colonization-specific isolators or individually ventilated cages. Germ-free Swiss Webster mice were orally gavaged with 13SM or 14SM at 6–8 weeks of age. Gnotobiotic mice and germ-free (GF) controls were fed the respective diets for at least 20 days prior to start of breeding. There was no significant difference in litter sizes between the 5 groups (13SM FR, 14SM FR, 14SM FF, GF FR, GF FF), however we did observe transient differences in pup weights according to diet and colonisation during weaning (**Fig EV3**). At 3 weeks of age, pups were weaned (i.e. physically separated from dams) and provided the same diet as the dams. Blinding was not possible as the colonization dictated the order of mouse handling and the diets are visually distinct.

### Diet formulation and administration

All parental mice were maintained on the FR diet, which is a standard rodent chow (SAFE®R04, Augy, France), until 2 weeks after the oral gavage. After this period, the parental mice designated as FF were switched to the custom fiber-free chow, which is a modified version of Harlan TD.08810 diet and is manufactured by the same vendor as the FR diet, as previously reported (Parrish, Grant, *et al*., 2022). Mice were maintained on the FF diet for at least 20 days prior to setting breeding pairs and the respective diets were administered *ad libitum* for the remainder of the study.

### Bacterial culturing and colonization of 13- or 14-member synthetic microbiota (SM)

Culturing and colonization of GF mice with the 13- or 14-member synthetic microbiota (SM) was performed as previously described (Steimle *et al*., 2021). The 14SM community contains the following bacteria (all type strains, unless otherwise indicated): *Akkermansia muciniphila*, *Collinsella aerofaciens*, *Desulfovibrio piger*, *Escherichia coli* HS, *Faecalibacterium prausnitzii* A2-165, *Roseburia intestinalis*, *Marvinbryantia formatexigens*, *Clostridium symbiosum*, *Eubacterium rectale* A1-86, *Barnesiella intestinihominis*, *Bacteroides caccae*, *Bacteroides uniformis*, *Bacteroides ovatus*, and *Bacteroides thetaiotaomicron*. The 13SM mice were orally gavaged with the same strains with the exception of *A. muciniphila*. Bacterial colonization among the parental mice was verified by qPCR of fecal DNA at 7 days post-oral gavage using strain-specific primers. Note that *R. intestinalis* and *E. rectale* did not consistently transfer to the pups during the early life period, despite being detected in the parents.

### Tissue processing

Distal colons were washed in PBS to collect contents, then transferred to 1 ml RNAprotect Tissue Reagent (Qiagen, Hilden, Germany) for colonic transcript analyses. Tissues were held in RNAprotect at 4°C for 2 days, after which the reagent was removed and the tissues stored at –80°C. For mice that would undergo FACS analyses, the entire colon was transferred into ice-cold 3 ml Hank’s balanced salt solution (HBSS) containing 10 mM HEPES and without Ca^2+^ and Mg^2+^. Cells from the colonic lamina propria (cLP) were isolated using a Lamina Propria Dissociation Kit (Miltenyi Biotec, Bergisch Gladbach, Germany) on the gentleMACS Dissociator (Miltenyi Biotec), following the manufacturer’s instructions. Cells were washed, counted, and re-suspended in FACS buffer (PBS; 1% FCS; 5mM EDTA) until cell staining.

### Flow cytometry

A total of 1.5 × 10^6^ cells per animal were transferred into a U-bottom 96-well plate for staining. For live/dead staining, cells were incubated in 100 µl PBS 1X containing 1.5 μL Zombie NIR Fixable Viability kit (BioLegend, San Diego, CA, USA) at 4°C for 30 min. Cells were washed twice using FACS buffer and subsequently fixed with the FoxP3 Fix/Perm kit (eBiosciences, Uithoorn, Netherlands) for 45 min at 4°C, followed by permeabilization wash. Purified Rat Anti-Mouse CD16/CD32 (Mouse BD Fc Block™, BD Biosciences, San Jose, CA, USA) was added at a concentration of 1 µg per 10^6^ cells, incubated at 4°C for 30 min, then cells were washed twice in permeabilization buffer. Cells were stained with anti-CD4 BV605 (#100548, clone RM4-5, 1:700, Biolegend), anti-CD3 BV711 (#100241, clone 17A2, 1:88, Biolegend), anti-CD45 BV780 (#564225, clone 30-F11, 1:88, BD), anti-CD335/NKp46 FITC (#137606, clone 29A1.4, 1:100, Biolegend), anti-CD8 PE-Cy5 (#100710, clone 53-6.7, 1:700, Biolegend), anti-FoxP3 eF450 (#48-5773-82, clone FJK-16s, 1:200, eBiosciences), anti-GATA3 PE (#100710, clone 53-6.7, 1:44, Biolegend), anti-EOMES PE-eF610 (#61-4875-82, clone Dan11mag, 1:100, eBiosciences), anti-Tbet PE-Cy7 (#644824, clone 4B10, 1:44, Biolegend), and anti-RORgt APC (#17-6988-82, clone AFKJS-9, 1:22, eBiosciences). Cells were incubated with the staining mix for 30 min at 4°C, then washed twice with FACS buffer. Samples were acquired on a NovoCyte Quanteon flow cytometer (ACEA Biosciences Inc., San Diego, CA, USA). Raw fcs files were analyzed in FlowJo version 10.8.1, with innate cell populations gated as described by Burrows et al. (Burrows *et al*., 2020). An example of the gating strategy is illustrated in **Fig EV2**. The dataset was exported as a percent of the total CD45^+^ cells for subsequent statistical analyses.

### DNA extraction and microbiome compositional analyses

Distal colon contents were collected by opening the colon longitudinally and washing tissue in 1 ml PBS. The PBS and contents were then centrifuged at 10 000 g for 10 min and the PBS supernatant was removed. Pelleted contents were stored at –20°C until DNA extraction using the phenol-chloroform method, as described by Steimle et al. (Steimle *et al*., 2021). The concentration of DNA was measured using a NanoPhotometer N60 (Implen, Munich, Germany). The purified DNA was diluted to 20 ng/ul and then subjected to qPCR with primers specific to each of the 14 bacterial strains, as previously described (Steimle *et al*., 2021). As the primers are specific to each strain and the target sequence is present only once in the respective genome (Desai *et al*., 2016), this method was used to allow for estimation of the number of each individual bacterial genomes or the absolute abundance of bacteria in the contents of the distal colon (**Dataset EV1–2**). As tissues were washed in 1X PBS to collect the contents, this estimation is based on the total contents, not normalized by the content weight, and can therefore vary according to factors such as mouse age, diet, and recent defecation. Streamplots based on the average abundance at each time point were created in Microsoft Excel 2016. In order to rule out contamination with non-14SM strains, we also performed 16S sequencing on a subset of samples. For this method, DNA concentration was assessed using the Qubit® dsDNA HS assay kit on a Qubit® 3.0 fluorometer (Life Technologies, Eugene, Oregon, USA). To rule out contamination with non-14SM members, sequencing of the V4 region of the 16S rRNA gene was carried out on an Illumina MiSeq system at the Integrated BioBank of Luxembourg (IBBL, Dudelange, Luxembourg), as previously described by Neumann et al. (Neumann *et al*., 2021). Note that 16S rRNA sequencing data is not well suited for analysis of 14SM because it cannot distinguish the three *Bacteroides* species well and because the variations in copy number can lead to biases in relative abundance calculations, nonetheless the results from the two methods are well-correlated for most of the SM strains (**Fig EV4**). Raw fastq sequences were processed via QIIME2 version 2020.6 (Bolyen *et al*., 2019) using DADA2 (Callahan *et al*., 2016) for quality control. Taxonomic assignment was carried out with the SILVA 138 database (Quast *et al*., 2013) (**Dataset EV3**).

### RNA extraction

Distal colon tissues were stored at –80°C until RNA extraction. Tissues were incubated overnight at –80°C in 1 ml TRIzol™ Reagent (Life Technologies Europe BV, Bleiswijk, Netherlands). A 5mm autoclaved metal bead was added to the sample, followed by homogenization on a RETSCH Mixer Mill MM 400 for 5 min at 30 Hz. Samples were centrifuged at 4°C for 5 min at 12 000 × g. The supernatant was transferred to a new tube and incubated for 5 min at room temperature. For each sample, 200 µl pure chloroform was added, then mixed by shaking for 15 s and incubated at room temperature for 3 min. Next, samples were centrifuged for 15 min at 4°C, 12 000 × g, and the aqueous phase was recovered. A volume of 500 µl isopropanol was added, followed by vigorous mixing and incubation at room temperature for 10 min. Samples were centrifuged at the same settings as previously. Then, the pellet was recovered and washed in 1 ml ice-cold 75% ethanol and vortexed briefly before centrifugation at 4°C for 5 min at 7 500 × g. The supernatant was discarded and the pellets were air-dried for 10 min at 37°C before re-suspending in 50 µl Invitrogen™ UltraPure™ DNase/RNase-free distilled water (Life Technologies Europe BV, Bleiswijk, Netherlands). The RNA solution was further incubated for 15 min at 56°C to fully re-suspend the pellet. DNase treatment to remove genomic DNA was done by adding 2 µl DNAse I (1 U/µl), 10 µl DNase I Buffer (10X), plus 40 µl water for 30 min at 37°C. Next, 1 µl of 0.5M EDTA was added to each sample and the DNAse I was inactivated by heating at 65°C for 10 min. RNA purification was performed with the RNeasy mini kit (QIAGEN Benelux BV, Venlo, Netherlands) according to manufacturer’s instructions. RNA was eluted in Invitrogen™ UltraPure™ DNase/RNase-free distilled water (Life Technologies Europe BV, Bleiswijk, Netherlands) and stored at –80°C.

### RT-qPCR

Complementary DNA libraries were prepared from 200 ng purified RNA in 11 µl water, 1 µl 2.5 uM random primers and µl 500 uM dNTPs (Life Technologies Europe BV, Bleiswijk, Netherlands), heated to 65°C for 5 min (lid temperature 112°C), then incubated at 4°C for at least 1 min. Afterwards, 1x SSIV buffer, 1 µl DTT, 1 uL RNaseOUT, and 1 uL SuperScript™ IV Reverse Transcriptase (Life Technologies Europe BV, Bleiswijk, Netherlands) was added to each tube. Samples were held at 23°C for 10 min, then 50°C for 10 min, and finally heated to 80°C for 10 min to inactivate enzymes. Quantitative PCR was carried out according to standard protocols, briefly: 1 ul of template cDNA was mixed with 8.7 µl water,1.2 µl 1x buffer, 0.625 µl 2.5 mM MgCl2, 0.5 µl 400uM dNTPs, 0.1 µl each of 0.2 uM forward and reverse primers (**Table 1**), 0.125 µl 1x SYBR green fluorescent dye, and 0.1 µl 0.5 U Taq DNA polymerase (Life Technologies Europe BV, Bleiswijk, Netherlands). Samples were pre-denatured at 94°C for 5 min; then underwent 40 cycles at 94°C for 20 s, 60°C for 50 s, 72°C for 45 s. Finally, samples were maintained at 72°C for 5 min and the melting curve was generated over 15 s by heating from 65°C–95°C at 0.3°C increments. Colonic transcript expression levels were normalized to levels of *Hprt*, which was identified as the most stable housekeeping gene over time and conditions using RefFinder (http://blooge.cn/RefFinder), in comparison to *Gapdh* and *Hsp90* (**Dataset EV8**).

**Table 1.**
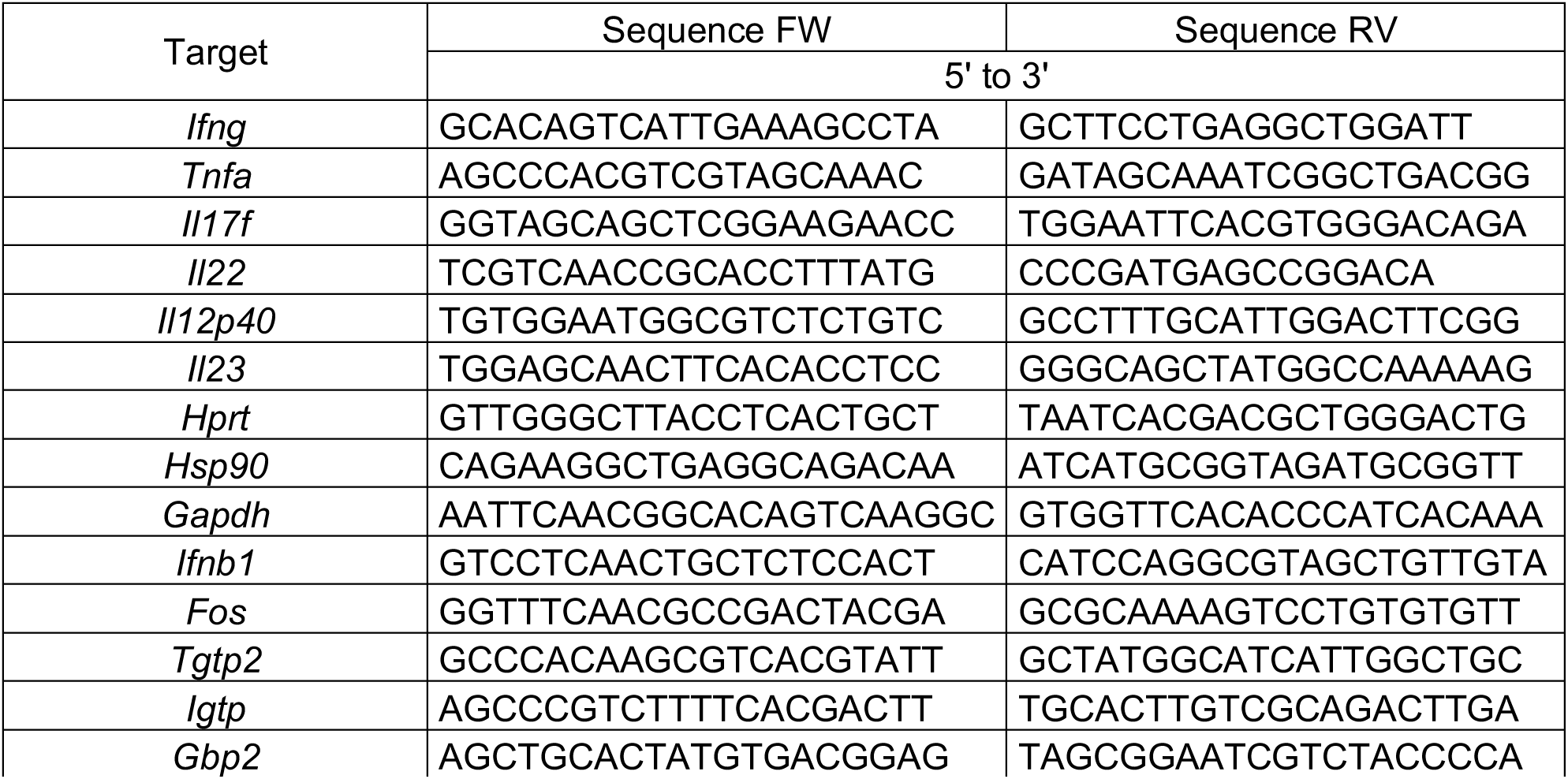
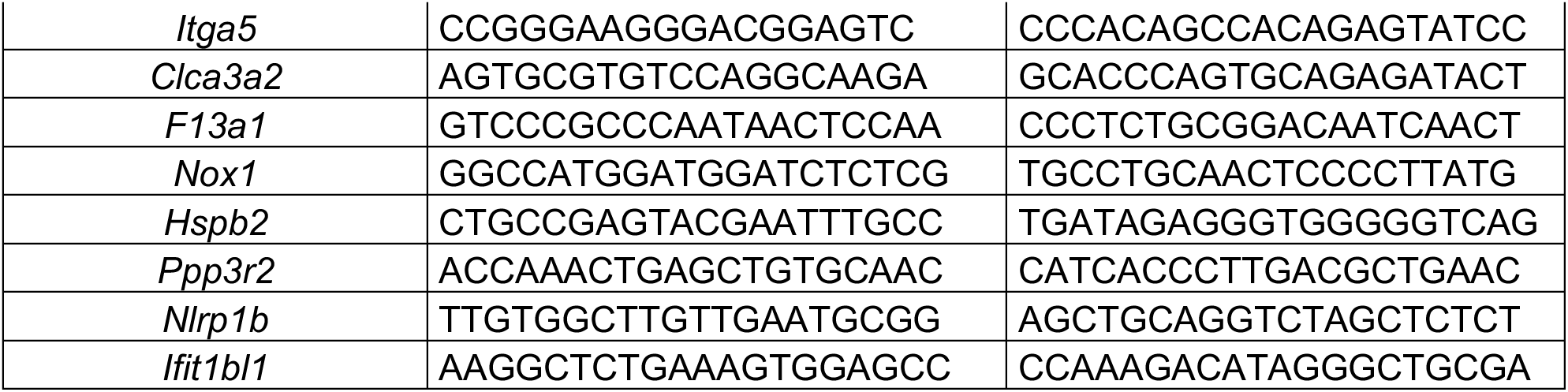
Primers for RT-qPCR against colonic cDNA transcript library.

### RNA sequencing and analysis

RNA integrity (RIN) was determined using a 2100 Bioanalyzer RNA Nanochip (Agilent, Santa Clara, CA, USA). Samples selected for RNA sequencing had RIN values ranging from 7.7–9.8. RNA libraries were prepared using an Illumina® Stranded Total RNA Prep kit, along with Ribo-Zero Plus to deplete ribosomal RNA. Samples were then sequenced in a 2 × 75 bp arrangement with a High Output Flow Cell on an Illumina NextSeq 550 system (Illumina, San Diego, CA, USA) at the LuxGen sequencing platform (Dudelange, Luxembourg). Adapters were removed using Cutadapt (Martin, 2011), followed by reads mapping and enumeration of gene counts with STAR 2.7.9a (Dobin *et al*., 2013) (**Dataset EV4**). Transcripts that did not appear at least one time on average across all samples were filtered from downstream analyses. Two outliers were identified based on variation on PC1 of the top 500 most variable genes and excluded from subsequent analyses (**Fig EV1A**). Samples normalization was performed using the median of ratios method, followed by differential expression analysis in R using DESeq2 1.30.1 (Love *et al*., 2014). Default parameters were used and *p* values were adjusted according to the Benjamini-Hochberg method. Differential expression was considered significant based on an adjusted *p* value<0.05. Differential expression analysis was performed for pairwise comparisons 14SM FF vs 14SM FR (**Dataset EV5**) and 14SM FR vs 13SM FR (**Dataset EV6**) (Love *et al*., 2014). Gene set analysis was also carried out in R using the clusterProfiler 3.18.1 enrichGO function (Yu *et al*., 2012) to identify up-or downregulated gene sets according to Gene Ontology (GO) terms (**Dataset EV7**).

### Statistical analysis

Unless otherwise specified, statistical analyses were carried out using GraphPad Prism version 9.3.1 for Windows (GraphPad Software, San Diego, CA, USA). For mRNA expression data by RT-qPCR, values were log-transformed prior to statistical analyses by one-way ANOVA and residuals tested for normality by the Kolmogorov-Smirnov test. For FACS data, outliers were identified in GraphPad using the ROUT method (Q = 10%), followed by a one-way ANOVA was used with multiple comparison correction using the Benjamini-Hochberg method. Only biologically-relevant pairwise statistical comparisons are shown on the plots: 13SM FR vs. 14SM FR, 13SM FR vs. GF FR, 14SM FR vs. 14SM FF, 14SM FR vs. GF FR, 14SM FF vs. GF FF, and GF FR vs. GF FF. Data are from independent experiments (pups born from up to 4 litters derived from 2–3 breeding pairs per condition) and represent biological replicates.

## Data Availability

The datasets produced in this study are available in the following databases:

- RNA-Seq and 16S rDNA data: European Nucleotide Archive (ENA) at EMBL-EBI PRJEB55622 (https://www.ebi.ac.uk/ena/browser/view/PRJEB55622)
- Flow cytometry data: FlowRepository (Spidlen *et al*., 2012) FR-FCM-Z5X6 (https://flowrepository.org/id/FR-FCM-Z5X6)

## Acknowledgements

Figure 4 was made using BioRender (Biorender.com). We thank the Luxembourg National Research Fund (FNR) for funding this research through CORE grants (C15/BM/10318186 and C18/BM/12585940) to M.S.D.; E.T.G. was supported by the FNR PRIDE (17/11823097) and the Fondation du Pélican de Mie et Pierre Hippert-Faber, under the aegis of the Fondation de Luxembourg; M.B. was supported by a European Commission Horizon 2020 Marie Skłodowska-Curie Actions individual fellowship (897408). We acknowledge the National Cytometry Platform (NCP) for assistance with generating flow cytometry data. The NCP is supported by Luxembourg’s Ministry of Higher Education and Research (MESR) funding.

## Author contributions

**Erica T Grant**: Conceptualization, data curation, formal analysis, investigation, methodology, software, validation, visualization, writing – original draft, writing – review & editing. **Marie Boudaud**: Conceptualization, data curation, formal analysis, investigation, methodology, validation, visualization, writing – review & editing. **Arnaud Muller**: Data curation, formal analysis, software, writing – review & editing. **Andrew J Macpherson**: Resources, writing – review & editing. **Mahesh S Desai**: Conceptualization, funding acquisition, investigation, methodology, project administration, resources, supervision, writing – review & editing

## Conflict of interest

Mahesh S. Desai works as a consultant and an advisory board member at Theralution GmbH, Germany.

## The Paper Explained

### PROBLEM

There is a growing awareness that the early life period is a critical window for shaping the course of neonatal development and future health outcomes. However, little is known regarding the effects of maternal diet and microbiome composition on the interconnected establishment of the gut microbiome and host immunity in the progeny. We sought to understand the role of maternal dietary fiber intake on early life development by breeding mice colonized with a 14-member synthetic microbiota (SM), fed a fiber-deficient diet. We further investigated the effects of discrete changes in the maternal microbiome by colonizing a third breeding pair with the full SM except *Akkermansia muciniphila*, a somewhat controversial, health-associated commensal that can digest milk oligosaccharides and host mucins.

## RESULTS

Pups born to dams fed a standard, fiber-rich chow showed initial colonization with *A. muciniphila*, followed by an expansion of fiber-degrading bacteria upon switching to solid food. Conversely, among pups of dams fed a fiber-free diet, *A. muciniphila* did not proliferate until after weaning, suggesting that maternal milk factors influenced by diet shape pups’ microbiome development. Pups of fiber-deprived dams expressed a significantly higher quantity of colonic tissue transcripts corresponding to defense response pathways against external antigens, and harbored lower proportions of group 2 innate lymphoid cells and Th17 cells. Pups of fiber-rich-fed mice lacking *A. muciniphila* demonstrated reduced proportions of innate and adaptive RORγt-positive immune cell subsets.

### IMPACT

The link between *A. muciniphila* and RORγt-positive immune cell subsets represents an intriguing potential to alter immunity by targeting a single bacterium, however the mechanisms underlying this interaction warrant further investigation. The unexpected suppression of *Akkermansia* among pups of fiber-free fed dams during the weaning period encourages the characterization of maternal milk components that might be used to support or limit growth of certain bacteria during early life. Using a tractable gnotobiotic mouse model and custom diet, we highlight the powerful potential to manipulate the microbiome and immunity during early life, which may have important implications for prevention of modern diseases such as allergy and autoimmunity.

## For more information

Gene Ontology (GO) Resource: http://geneontology.org/

RefFinder: http://blooge.cn/RefFinder/

Eco-Immunology and Microbiome Group (EIM): https://eim.lih.lu

Luxembourg Genome Center (LuxGen): https://luxgen.lih.lu

Luxembourg Institute of Health Research Portal: https://researchportal.lih.lu/

## Expanded View

**Figure EV1.**
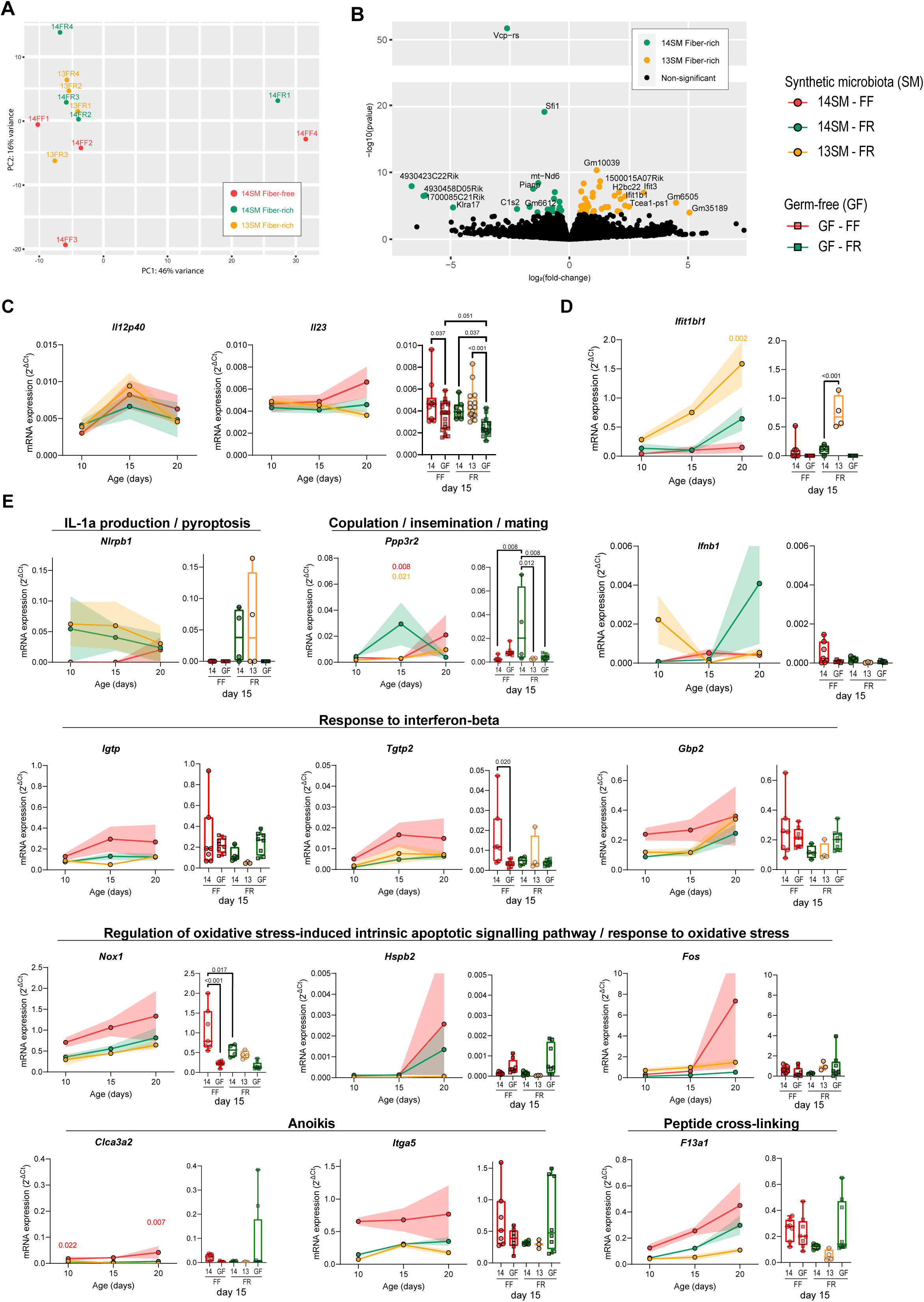
**A**. PCA plot of the top 500 most variable genes among samples analyzed by RNA sequencing (*n* = 4 mice/group). **B**. Volcano plot showing colonic transcripts that were significantly enriched in 14SM FR or 13SM FR after correction for multiple comparisons using DESeq2 (*n* = 3-4 mice/group). **C**. mRNA expression of *Il12p40* and *Il23*, targeted transcripts of interest (*n* = 17-24 mice/group). **D**. mRNA expression of *Ifit1b1*, transcript identified by RNA sequencing as elevated in 13SM FR. **E**. mRNA expression of transcripts belonging to top 5 GO gene sets elevated in 14SM FR (*Nlrbp1* and *Ppp3r2*) or 14SM FF (*Igtp*, *Tgtp2*, *Gbp2*, *Nox1*, *Hspb2*, *Fos*, *Clca3a2*, *Itga5*, *a*nd *F13a1*). Colonic transcription levels were normalized by *Hprt*, which was the most stable among the three housekeeping genes tested (*Hprt, Hsp90, Gapdh)*. Data information: Longitudinal data of mean ± SEM is shown for 14SM FF, 14SM FR, and 13SM FR pups at age 10-20 days (circles) with statistical significance calculated using two-way ANOVA factored on group and time point compared to the 14SM FR control. To the right of each longitudinal plot, we also report colonic transcript levels for pups born to germ-free (GF) FF or FR dams at age 15 days (squares) with statistical significance based on a one-way ANOVA with false-discovery adjustment using the Benjamini and Hochberg method. Outliers removed using ROUT method with Q = 10%.

**Figure EV2.**
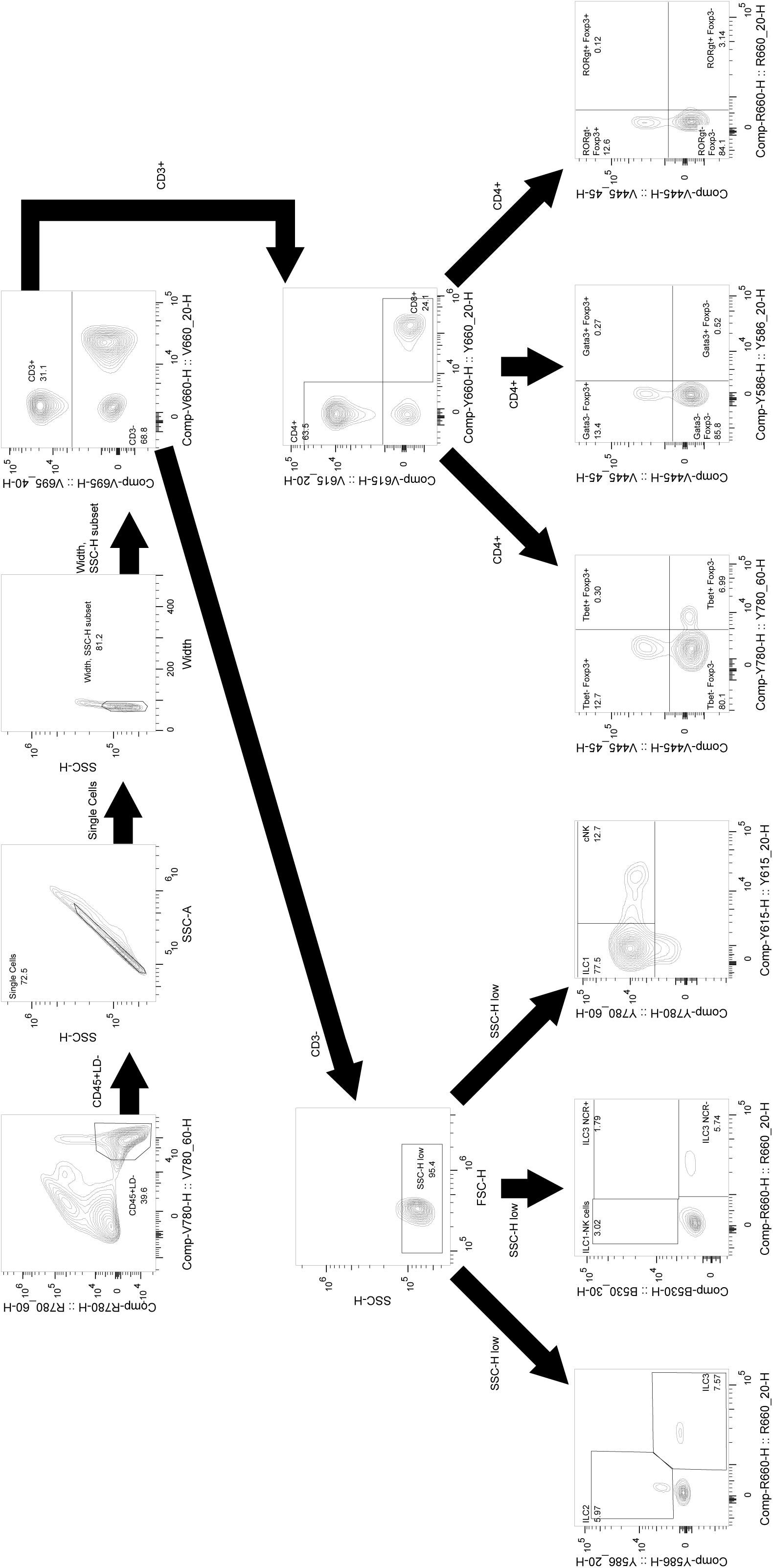
Representative gating strategy for flow cytometry analysis.

**Figure EV3.**
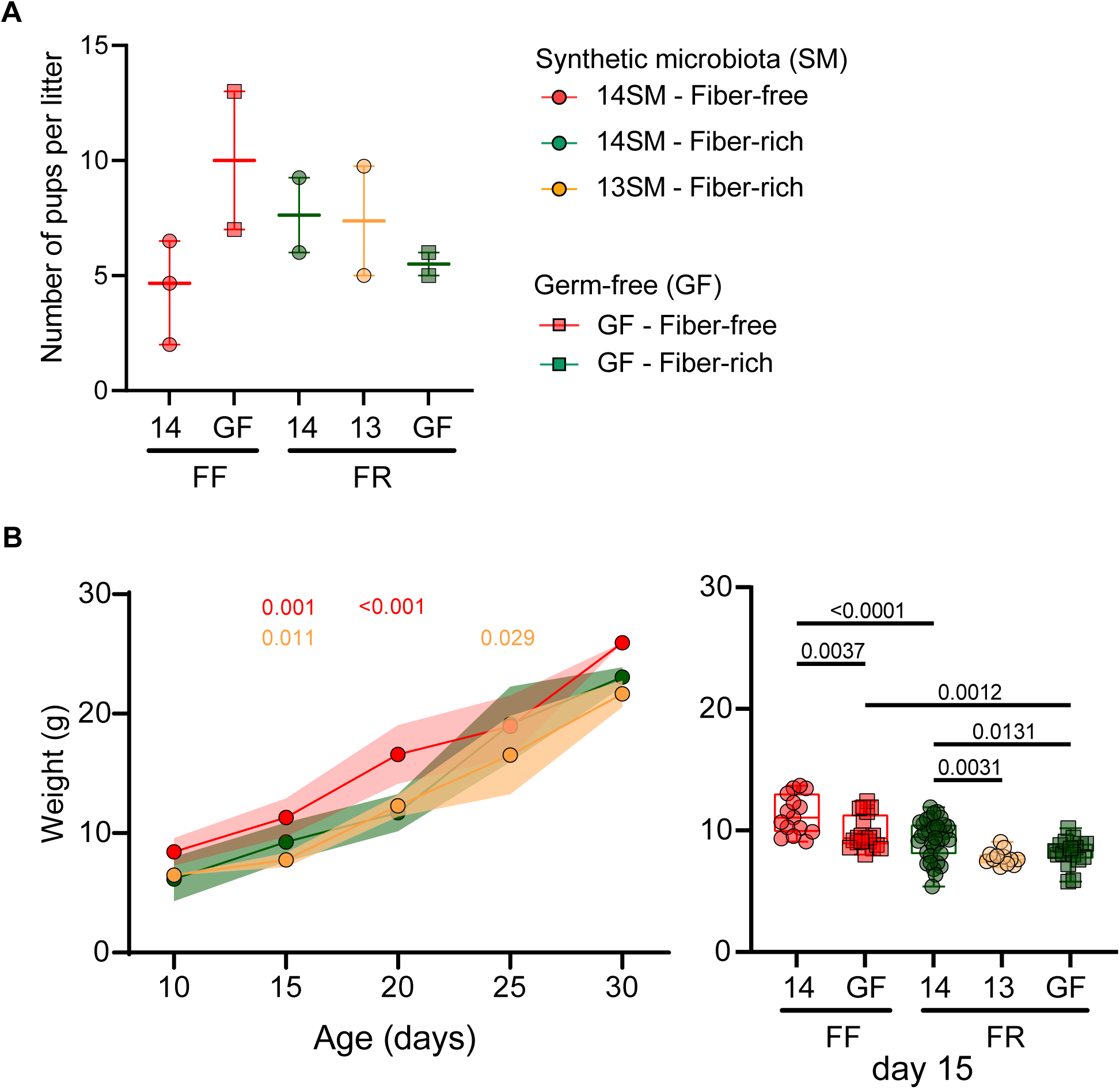
**A**. Number of pups born per litter in each group. (*n* = 2-3 litters/group). No statistical significance for pairwise comparisons based on a one-way ANOVA with false-discovery adjustment using the Benjamini and Hochberg method. **B**. Pup weights at the specified intervals. (*n* = 2-17 mice/group). Longitudinal data of mean weight (g) ± SEM is shown for 14SM FF, 14SM FR, and 13SM FR pups at age 10-30 days (circles) with statistical significance calculated using two-way ANOVA factored on group and time point compared to the 14SM FR control. To the right, we also report weights of pups born to germ-free (GF) FF or FR dams at age 15 days (squares) with statistical significance based on a one-way ANOVA with false-discovery adjustment using the Benjamini and Hochberg method.

**Figure EV4.**
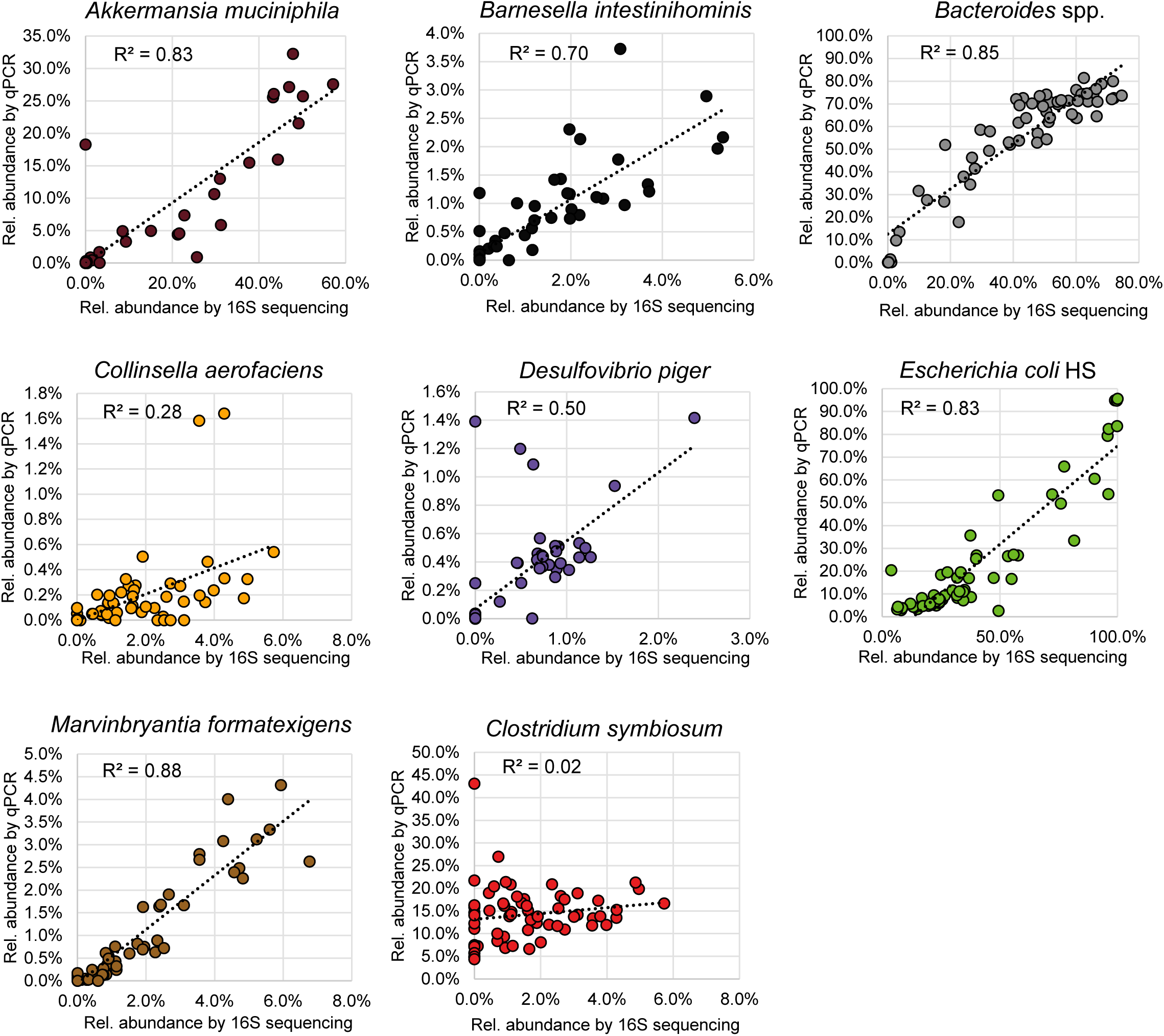
Correlation coefficient (R^2^) for the linear trendline fit to relative abundance by qPCR (y- axis) versus relative abundance by 16S rRNA gene sequencing (x-axis). Plots shown for all bacteria that were present at a detectable level in the pups of dams colonized with 13SM or 14SM.

**Dataset EV1.** Abundances of synthetic microbiota (SM) members among parent mice by qPCR. Compositions are based on fecal samples taken at the end of the study.

**Dataset EV2.** Abundances of synthetic microbiota (SM) members among individual pups by qPCR.

**Dataset EV3.** 16S rRNA gene sequencing to rule out contamination with non-14SM bacteria.

**Dataset EV4.** Count table of colonic transcripts mapping to genes by ENSEMBL identifier. Asterisk (*) indicates samples identified as outliers by PCA plotting (**Fig EV1A**).

**Dataset EV5.** Differential enrichment analysis using DESeq2 for 14SM fiber-free and 14SM fiber- rich contrasts.

**Dataset EV6.** Differential enrichment analysis using DESeq2 for 14SM fiber-rich and 13SM fiber- rich contrasts.

**Dataset EV7.** Gene set differential enrichment analysis using clusterProfiler (biological processes) for 14SM fiber-free and 14SM fiber-rich contrasts.

**Dataset EV8.** RefFinder results to identify most stable housekeeping gene.

## Notes

### Summary of Updates

The manuscript has been updated as per the reviewer comments. We have carefully discussed the points raised by the reviewers, which we have found to be highly useful in elaborating on the discussion of our findings and future directions. We have been able to address the reviewers concerns by performing new experiments (longitudinal RT-qPCR in Figure EV1); re-analyzing the existing data to address questions over outliers, data transformations and multiple comparison corrections; and by providing additional discussion of our results, where requested. In our response to reviewer comments, we explain the reasons that we were unable to perform metabolomics analysis on existing samples. We have also added a few lines to the manuscript to discuss the relevance of metabolomics analysis in our model and predictions of potential changes that would be important to verify in follow-up studies: Regarding the timing of the diet switch and potential cross-fostering experiments, we have: 1) explained the rationale for not incorporating additional animal work into the existing manuscript in our response to the reviewers 2) clarified important aspects of the study design in the Materials and Methods sections Mice and Diet formulation and administration and 3) added discussion of follow-up points to investigate in future studies in the manuscript.

## References

1. Afrizal A, Jennings SA V, Hitch TCA, Riedel T, Basic M, et al. (2022) Enhanced cultured diversity of the mouse gut microbiota enables custom-made synthetic communities. Cell Host Microbe 30 (11), 1630–1645.

2. De Agüero MG, Ganal-Vonarburg SC, Fuhrer T, Rupp S, Uchimura Y, et al. (2016) The maternal microbiota drives early postnatal innate immune development. Science. 351 (6279), 1296– 1302.

3. Ansaldo E, Slayden LC, Ching KL, Koch MA, Wolf NK, et al. (2019) Akkermansia muciniphila induces intestinal adaptive immune responses during homeostasis. Science. 364 (6446), 1179–1184.

4. Arrigo A-P, Simon S, Gibert B, Kretz-Remy C, Nivon M, et al. (2007) Hsp27 (HspB1) and αB-crystallin (HspB5) as therapeutic targets. FEBS Lett. 581 (19), 3665–3674.

5. Babu ST, Niu X, Raetz M, Savani RC, Hooper L V, et al. (2018) Maternal high-fat diet results in microbiota-dependent expansion of ILC3s in mice offspring. JCI insight 3 (19)

6. Blachier F, Andriamihaja M, Larraufie P, Ahn E, Lan A, et al. (2021) Production of hydrogen sulfide by the intestinal microbiota and epithelial cells and consequences for the colonic and rectal mucosa. Am. J. Physiol. Liver Physiol. 320 (2), G125–G135.

7. Bolyen E, Rideout JR, Dillon MR, Bokulich NA, Abnet CC, et al. (2019) Reproducible, interactive, scalable and extensible microbiome data science using QIIME 2. Nat. Biotechnol.

8. Brown RL, Larkinson MLY, and Clarke TB (2021) Immunological design of commensal communities to treat intestinal infection and inflammation. PLoS Pathog. 17 (1), e1009191.

9. Burrows K, Chiaranunt P, Ngai L, and Mortha A (2020) Rapid isolation of mouse ILCs from murine intestinal tissues. In: Methods in Enzymology. Elsevier, Vol 631, pp. 305–327

10. Callahan BJ, McMurdie PJ, Rosen MJ, Han AW, Johnson AJA, et al. (2016) DADA2: High-resolution sample inference from Illumina amplicon data. Nat. Methods 13 (7), 581–583.

11. Cheng D and Xie MZ (2021) A review of a potential and promising probiotic candidate—Akkermansia muciniphila. J. Appl. Microbiol. 130 (6), 1813–1822.

12. Cirstea M, Radisavljevic N, and Finlay BB (2018) Good bug, bad bug: breaking through microbial stereotypes. Cell Host Microbe 23 (1), 10–13.

13. Collado MC, Derrien M, Isolauri E, de Vos WM, and Salminen S (2007) Intestinal integrity and Akkermansia muciniphila, a mucin-degrading member of the intestinal microbiota present in infants, adults, and the elderly. Appl. Environ. Microbiol. 73 (23), 7767–7770.

14. Davis EC, Castagna VP, Sela DA, Hillard MA, Lindberg S, et al. (2022) Gut microbiome and breast-feeding: Implications for early immune development. J. Allergy Clin. Immunol. 150 (3), 523– 534.

15. Desai MS, Seekatz AM, Koropatkin NM, Kamada N, Hickey CA, et al. (2016) A Dietary Fiber-Deprived Gut Microbiota Degrades the Colonic Mucus Barrier and Enhances Pathogen Susceptibility. Cell 167 (5), 1339–1353.e21.

16. Dobin A, Davis CA, Schlesinger F, Drenkow J, Zaleski C, et al. (2013) STAR: ultrafast universal RNA-seq aligner. Bioinformatics 29 (1), 15–21.

17. Earle KA, Billings G, Sigal M, Lichtman JS, Hansson GC, et al. (2015) Quantitative Imaging of Gut Microbiota Spatial Organization. Cell Host Microbe 18 (4), 478–488.

18. Ganal SC, Sanos SL, Kallfass C, Oberle K, Johner C, et al. (2012) Priming of natural killer cells by nonmucosal mononuclear phagocytes requires instructive signals from commensal microbiota. Immunity 37 (1), 171–186.

19. Geerlings SY, Kostopoulos I, De Vos WM, and Belzer C (2018) Akkermansia muciniphila in the human gastrointestinal tract: when, where, and how? Microorganisms 6 (3), 75.

20. Gu Z, Pei W, Shen Y, Wang L, Zhu J, et al. (2021) Akkermansia muciniphila and its outer protein Amuc_1100 regulates tryptophan metabolism in colitis. Food Funct. 12 (20), 10184–10195.

21. Hayase E, Hayase T, Jamal MA, Miyama T, Chang C-C, et al. (2022) Mucus-degrading Bacteroides link carbapenems to aggravated graft-versus-host disease. Cell 185 (20), 3705–3719.

22. Hepworth MR, Fung TC, Masur SH, Kelsen JR, McConnell FM, et al. (2015) Group 3 innate lymphoid cells mediate intestinal selection of commensal bacteria–specific CD4+ T cells. Science. 348 (6238), 1031–1035.

23. Horvath TD, Ihekweazu FD, Haidacher SJ, Ruan W, Engevik KA, et al. (2022) Bacteroides ovatus colonization influences the abundance of intestinal short chain fatty acids and neurotransmitters. Iscience 25 (5), 104158.

24. Hrncir T, Stepankova R, Kozakova H, Hudcovic T, and Tlaskalova-Hogenova H (2008) Gut microbiota and lipopolysaccharide content of the diet influence development of regulatory T cells: studies in germ-free mice. BMC Immunol. 9 (1), 1–11.

25. Jangi S, Gandhi R, Cox LM, Li N, von Glehn F, et al. (2016) Alterations of the human gut microbiome in multiple sclerosis. Nat. Commun. 7 (1), 12015.

26. Kalbermatter C, Fernandez Trigo N, Christensen S, and Ganal-Vonarburg SC (2021) Maternal Microbiota, Early Life Colonization and Breast Milk Drive Immune Development in the Newborn. Front. Immunol. 12, 1768.

27. Kennedy EA, King KY, and Baldridge MT (2018) Mouse microbiota models: comparing germ-free mice and antibiotics treatment as tools for modifying gut bacteria. Front. Physiol. 9, 1534.

28. Kostopoulos I, Elzinga J, Ottman N, Klievink JT, Blijenberg B, et al. (2020) Akkermansia muciniphila uses human milk oligosaccharides to thrive in the early life conditions in vitro. Sci. Rep. 10 (1), 1–17.

29. Li S, Bostick JW, Ye J, Qiu J, Zhang B, et al. (2018) Aryl hydrocarbon receptor signaling cell intrinsically inhibits intestinal group 2 innate lymphoid cell function. Immunity 49 (5), 915–928.

30. Lin X, Tawch S, Singh A, Morgun A, Shulzhenko N, et al. (2021) Akkermansia muciniphila induces Th17 cells by mediating tryptophan metabolism and exacerbates experimental autoimmune encephalomyelitis (EAE). J. Immunol. 206 (1 Supplement), 105.09 LP-105.09.

31. Liu Y, Song Y, Lin D, Lei L, Mei Y, et al. (2019) NCR− group 3 innate lymphoid cells orchestrate IL-23/IL-17 axis to promote hepatocellular carcinoma development. EBioMedicine 41, 333–344.

32. Liu Y, Yang M, Tang L, Wang F, Huang S, et al. (2022) TLR4 regulates RORγt+ regulatory T-cell responses and susceptibility to colon inflammation through interaction with Akkermansia muciniphila. Microbiome 10 (1), 1–20.

33. Love MI, Huber W, and Anders S (2014) Moderated estimation of fold change and dispersion for RNA-seq data with DESeq2. Genome Biol. 15 (12), 550.

34. Lubin J-B, Green J, Maddux S, Denu L, Duranova T, et al. (2023) Arresting microbiome development limits immune system maturation and resistance to infection in mice. Cell Host Microbe

35. Luna E, Parkar SG, Kirmiz N, Hartel S, Hearn E, et al. (2022) Utilization efficiency of human milk oligosaccharides by human-associated Akkermansia is strain dependent. Appl. Environ. Microbiol. 88 (1), e01487–21.

36. Macpherson AJ and Harris NL (2004) Interactions between commensal intestinal bacteria and the immune system. Nat. Rev. Immunol. 4 (6), 478–485.

37. Martens E, Pereira G, Boudaud M, Wolter M, Alexander C, et al. (2023) Unravelling specific diet and gut microbial contributions to inflammatory bowel disease. ResearchSquare https://doi.org/10.21203/rs.3.rs-2518251/v1 [PREPRINT]

38. Martin M (2011) Cutadapt removes adapter sequences from high-throughput sequencing reads. EMBnet. J. 17 (1), 10–12.

39. Mirpuri J (2021) Evidence for maternal diet-mediated effects on the offspring microbiome and immunity: implications for public health initiatives. Pediatr. Res. 89 (2), 301–306.

40. Al Nabhani Z, Dulauroy S, Marques R, Cousu C, Al Bounny S, et al. (2019) A Weaning Reaction to Microbiota Is Required for Resistance to Immunopathologies in the Adult. Immunity 50 (5), 1276–1288.

41. Neumann M, Steimle A, Grant ET, Wolter M, Parrish A, et al. (2021) Deprivation of dietary fiber in specific-pathogen-free mice promotes susceptibility to the intestinal mucosal pathogen Citrobacter rodentium. Gut Microbes

42. Parrish A, Boudaud M, Grant E, Willieme S, Neumann M, et al. (2022) Akkermansia muciniphila regulates food allergy in a diet-dependent manner. ResearchSquare https://doi.org/10.21203/rs.3.rs-1745691/v1 [PREPRINT]

43. Parrish A, Grant E, Boudaud M, Hunewald O, Hirayama A, et al. (2022) Dietary fibers boost gut microbiota-produced B vitamin pool and alter host immune landscape. ResearchSquare https://doi.org/doi.org/10.21203/rs.3.rs-1563674/v2 [PREPRINT]

44. Qiu J, Heller JJ, Guo X, Zong-ming EC, Fish K, et al. (2012) The aryl hydrocarbon receptor regulates gut immunity through modulation of innate lymphoid cells. Immunity 36 (1), 92–104.

45. Quast C, Pruesse E, Yilmaz P, Gerken J, Schweer T, et al. (2013) The SILVA ribosomal RNA gene database project: Improved data processing and web-based tools. Nucleic Acids Res.

46. Rey FE, Faith JJ, Bain J, Muehlbauer MJ, Stevens RD, et al. (2010) Dissecting the in vivo metabolic potential of two human gut acetogens. J. Biol. Chem. 285 (29), 22082–22090.

47. Robertson RC, Manges AR, Finlay BB, and Prendergast AJ (2019) The human microbiome and child growth–first 1000 days and beyond. Trends Microbiol. 27 (2), 131–147.

48. Sanos SL, Bui VL, Mortha A, Oberle K, Heners C, et al. (2009) RORγt and commensal microflora are required for the differentiation of mucosal interleukin 22–producing NKp46+ cells. Nat. Immunol. 10 (1), 83–91.

49. Sonnenburg ED, Smits SA, Tikhonov M, Higginbottom SK, Wingreen NS, et al. (2016) Diet-induced extinctions in the gut microbiota compound over generations. Nature 529 (7585), 212–215.

50. Spidlen J, Breuer K, Rosenberg C, Kotecha N, and Brinkman RR (2012) FlowRepository: A resource of annotated flow cytometry datasets associated with peer-reviewed publications. Cytom. Part A

51. Steimle A, De Sciscio A, Neumann M, Grant ET, Pereira G V., et al. (2021) Constructing a gnotobiotic mouse model with a synthetic human gut microbiome to study host–microbe cross talk. STAR Protoc. 2 (2)

52. Szanto I, Rubbia-Brandt L, Kiss P, Steger K, Banfi B, et al. (2005) Expression of NOX1, a superoxide-generating NADPH oxidase, in colon cancer and inflammatory bowel disease. J. Pathol. A J. Pathol. Soc. Gt. Britain Irel. 207 (2), 164–176.

53. Takeuchi T, Miyauchi E, Kanaya T, Kato T, Nakanishi Y, et al. (2021) Acetate differentially regulates IgA reactivity to commensal bacteria. Nature 595 (7868), 560–564.

54. Thorburn AN, McKenzie CI, Shen SJ, Stanley D, Macia L, et al. (2015) Evidence that asthma is a developmental origin disease influenced by maternal diet and bacterial metabolites. Nat. Commun. 6 (1), 7320.

55. Wells JM, Gao Y, de Groot N, Vonk MM, Ulfman L, et al. (2022) Babies, Bugs, and Barriers: Dietary Modulation of Intestinal Barrier Function in Early Life. Annu. Rev. Nutr. 42

56. Xie Y-H, Gao Q-Y, Cai G-X, Sun X-M, Zou T-H, et al. (2017) Fecal Clostridium symbiosum for noninvasive detection of early and advanced colorectal cancer: test and validation studies. EBioMedicine 25, 32–40.

57. Yoon HS, Cho CH, Yun MS, Jang SJ, You HJ, et al. (2021) Akkermansia muciniphila secretes a glucagon-like peptide-1-inducing protein that improves glucose homeostasis and ameliorates metabolic disease in mice. Nat. Microbiol. 6 (5), 563–573.

58. Yu G, Wang L-G, Han Y, and He Q-Y (2012) clusterProfiler: an R package for comparing biological themes among gene clusters. Omi. a J. Integr. Biol. 16 (5), 284–287.

59. Zheng W, Zhao W, Wu M, Song X, Caro F, et al. (2020) Microbiota-targeted maternal antibodies protect neonates from enteric infection. Nature 577 (7791), 543–548.

60. Zhu Hai, Wang G, Zhu Haixing, and Xu A (2021) ITGA5 is a prognostic biomarker and correlated with immune infiltration in gastrointestinal tumors. BMC Cancer 21 (1), 1–14.

